# Transcription factor AhR regulates glutathione S-transferases (GSTs) conferring resistance to *lambda*-cyhalothrin in *Cydia pomonella*

**DOI:** 10.1101/2023.01.03.522531

**Authors:** Chao Hu, Yu-Xi Liu, Shi-Pang Zhang, Ya-Qi Wang, Ping Gao, Yu-Ting Li, Xue-Qing Yang

## Abstract

Transcription factor aryl hydrocarbon receptor (AhR) can enhance insect resistance to insecticides by regulating the detoxification metabolic network. Our previous studies have confirmed that overexpression of cytochrome P450 monooxygenases (P450s) and glutathione S-transferases (GSTs) are both involved in *lambda*-cyhalothrin resistance in *Cydia pomonella*. In this study, we report that AhR regulates GSTs thus conferring *lambda*-cyhalothrin resistance in *C. pomonella*. Spatiotemporal expression patterns indicated that *AhR* gene of *C. pomonella* (*CpAhR*) was highly expressed in the Malpighian tubules of larvae. Moreover, the expression of *CpAhR* was induced by *lambda*-cyhalothrin exposure and was up-regulated in a *lambda*-cyhalothrin-resistant population. RNA interference (RNAi) of the expression of *CpAhR* could effectively decrease the relative expression level of *CpGSTe3* and enzyme activity of GSTs, but not P450s, further reducing the tolerance of larvae to *lambda*-cyhalothrin. Furthermore, β-naphthoflavone (BNF), a novel agonist of AhR, can effectively increase the expression of *CpAhR* and the activity of the GSTs enzyme, resulting in the enhancement of larvae tolerance to *lambda*-cyhalothrin. These results demonstrate that *lambda*-cyhalothrin exposure can effectively activate the expression of *CpAhR* and increase GSTs enzyme thus leading to the development of resistance to *lambda*-cyhalothrin, which enriches the theory of insecticide resistance regulation in *C. pomonella*.

## 1. Introduction

The codling moth, *Cydia pomonella* (L.) (Lepidoptera: Tortricidae), is a major pest on pome fruits that causes huge economic losses worldwide [1]. Most of the duration of the larval stage of *C. pomonella* occurs inside the fruit, making it difficult to be controlled [2]. Although environment-friendly control methods have been sought in agroforestry management, chemical insecticide is still the most effective measure [3,4]. *Lambda*-cyhalothrin is regarded as one of the most commonly used chemical insecticides to control *C. pomonella* [5]. However, insecticide overuse has led to the development of *C. pomonella* resistance to *lambda*-cyhalothrin [6,7]. Our previous study has shown that several GST genes were overexpressed in a field *lambda*-cyhalothrin-resistant population of *C. pomonella* [8]. Recently, the recombinant proteins of CpGSTd1, CpGSTd3, CpGSTe3, and CpGSTs2 were found to be able to metabolize *lambda*-cyhalothrin via sequestration, implying that GSTs may be involved in the resistance of *lambda*-cyhalothrin [8]. In addition, it is confirmed that the metabolic functional redundancy of three P450 genes in the *CYP9A* subfamily leads to P450-mediated *lambda*-cyhalothrin resistance in *C. pomonella* [9]. However, the transcriptional regulation mechanism of insecticide resistance genes is unclear in *C. pomonella*.

In insects, the regulation of the expression of detoxification genes can be carried out at multiple levels, and among them, transcriptional regulation has been regarded as the most important link of gene regulation, which is mainly affected by *trans*-acting factors (transcription factors) and *cis*-regulatory elements [10,11]. At present, transcription factors related to the regulation of detoxification genes and further conferring insecticide resistance are mainly distributed in three super-families: the nuclear receptor (NR) transcription factors super-family, the basic helix-loop-helix-Per-Arnt-Sim domain (bHLH-PAS) transcription factors super-family, and the basic leucine zipper (bZIP) transcription factors super-family [12]. The aryl hydrocarbon receptor (AhR), a member of the bHLH-PAS super-family, is a ligand-activated transcription factor, which participates in exerting both physiological stress and expression regulation to exogenous toxicants [12-14]. The expression of the *AhR* gene can be activated by a large number of exogenous or endogenous environmental factors [15,16]. It is transferred into the nucleus where it combines with another constitutive protein, the AhR nuclear translocator (ARNT), and the heterodimer of AhR/ARNT can recognize specific binding elements to regulate gene expression, including activation and inhibition [17,18]. This transcription factor is well conserved among mammals, fish, birds, and arthropods, which drives them to perform relatively conservative functions, including regulation of detoxification and metabolism of xenobiotics [19-21]. However, only a few studies have focused on the regulation of the AhR/ARNT regulatory pathway in the expression of phase I and phase II enzymes in the insecticide metabolic process in insects. It has been proved that *CYP6CY3* is transcriptionally regulated by *AhR* and *ARNT* in *Myzus persicae* and conferred an adaptive evolution to nicotine [22]. Knockdown of the expression of the *AhR* gene decreased the P450 enzyme activity and reduced the tolerance of *Nilaparvata lugens* to several insecticides [23]. The transcription factor AhR of *Dendroctonus armandi* was induced by terpenoids and participated in the regulation of *DaCYP6DF1* expression, which was associated with susceptibility to (−)-β-pinene and (±) -limonene [24]. Therefore, clarifying the regulatory relationship between the transcription factor AhR and detoxification enzymes is critical for better understanding resistance mechanisms to insecticides and provides important clues for improving sustainable pest management. However, the transcription factor AhR has not been identified in *C. pomonella* and its contribution to the regulation of detoxifying gene transcription is unknown.

Modulating activity has been considered to be an essential parameter of verifying the adaptability of transcription factors to exogenous stress [25], and the signaling is considered to be regulated at three levels as follows: proteasomal degradation of the AhR, AhR-ARNT complex disruption by AhR repressor (AHRR), and ligand metabolism by CYP enzymes [26]. Agonists and antagonists are commonly used to study the regulatory activity of transcription factors. However, there are few regulators used to investigate the transcription factor *AhR*, especially in insects. β-naphthoflavone (BNF) is a synthetic derivative of natural flavonoids, which has long been identified as a potent agonist [27], and it has been identified to induce the activity of a variety of phase I and phase II enzymes in mammals and fish [28,29]. However, in arthropods, there are no relevant studies to evaluate the activity of agonist BNF to stimulate the transcription factor *AhR*. In the present study, we identified and characterized the AhR from *C. pomonella* (*CpAhR*). We then examined whether the overexpression of detoxification enzymes is AhR-mediated. Our results can help in revealing the transcriptional regulation mechanism of insecticide resistance genes and are important for the resistance management of *C. pomonella*.

## 2. Materials and methods

### 2.1. Insects and chemicals

A susceptible strain (SS) of *C. pomonella* was maintained for more than 50 generations without exposure to insecticides [1]. A field-resistant strain (ZW_R, resistant ratio=16.97) and its resistant regression strain (ZW_S, resistant ratio=1.53) [8], were maintained in the laboratory by using the method of artificial breeding referred to Wang et al [5]. In short, The larvae were fed with artificial feed in an artificial incubator under a 16 h light: 8 h dark (L:D) cycle at 26 ± 1 ℃ and 60 ± 5 % humidity. The moths were transferred to a self-made storage box and fed with 10% honey diluted with pure water.

Technical-grade (purity >98%) *lambda*-cyhalothrin was obtained from Aladdin Reagent (Shanghai, China), β-naphthoflavone (BNF) (purity >99%), and dimethyl sulphoxide (DMSO) (purity =99.9%) were supplied by Beijing Solarbio Science & Technology Co., Ltd. (Beijing, China).

### 2.2. Insect exposures and sample preparation

The sublethal dose (LD_10_) or half-lethal dose (LD_50_) value of *lambda*-cyhalothrin [5] was treated to larvae for enzyme activity and bioassay analysis. One μL drop of suitable concentration of *lambda*-cyhalothrin was applied on the thoracic dorsum of each fourth-instar larva and each sample was collected at 6, 12, 24, and 48 hours post-exposure (hpe). To determine the role of agonist BNF in activating transcription factor AhR, we preselected several concentrations of BNF for the feeding experiment, which is based on the approximate conversion in other species [28]. In brief, BNF dissolved in DMSO was added to the artificial diet at 1.0, 1.5, and 2.0 mg g^-1^ respectively, and the same volume of DMSO was added as a control. The newly hatched fourth-instar larvae were transferred to the diet supplemented with different concentrations of BNF. After recording the weight changes within 0 h to 48 h, test larvae were collected for determination of enzyme activity and tolerance to *lambda*-cyhalothrin. A total of three groups were repeated and five and fifteen larvae were collected for determination of enzyme activities and insecticide tolerance for each repeat, respectively. For different developmental stages and tissues used in expression profiling, sufficient biological individual samples were selected, according to the requirements of RNA extraction instructions. All samples were rapidly frozen in liquid nitrogen and stored at - 80 ℃.

### 2.3. Total RNA extraction, cDNA synthesis, and RT-qPCR

The total RNA of all samples of *C. pomonella* was isolated using RNAiso Plus (Takara, Beijing, China) following the manufacturer’s instructions, and the concentration of extracted total RNA was estimated by NanoDrop 2000 (Thermo-Fisher Scientific). As needed, the first-strand cDNA was synthesized from 1 μg total RNA using PrimeScript™ II 1st Strand cDNA Synthesis Kit (Takara, Beijing, China) following the manufacturer’s instructions.

The mRNA levels were assessed by real-time quantitative polymerase chain reaction (RT-qPCR) with Bio-Rad CFX96 (Bio-Rad), and a 20 μL of reaction, containing 1 μL of proper concentration of cDNA template, 10 μL of TB Green Premix Ex Taq 2 (TliRNaseH Plus) (Takara, Beijing, China), 0.8 μL of 10 μM bidirectional primers (Table 1), and moderate water for constant volume, was premixed for PCR cycling condition as follows: initial denaturation at 95 ℃ for 30 s, followed by 40 cycles of 95 ℃ for 5 s and 60 ℃ for 30 s, and final melt curve analysis from 55 to 95 ℃. *EF-1α* (MN037793) and *GAPDH* (MT116773) [7] were used as internal control genes in RT-qPCR. Each experiment was performed with three biological replicates. The relative expression of each gene was calculated using the 2^-ΔΔCT^ method [30].

**Table 1.**
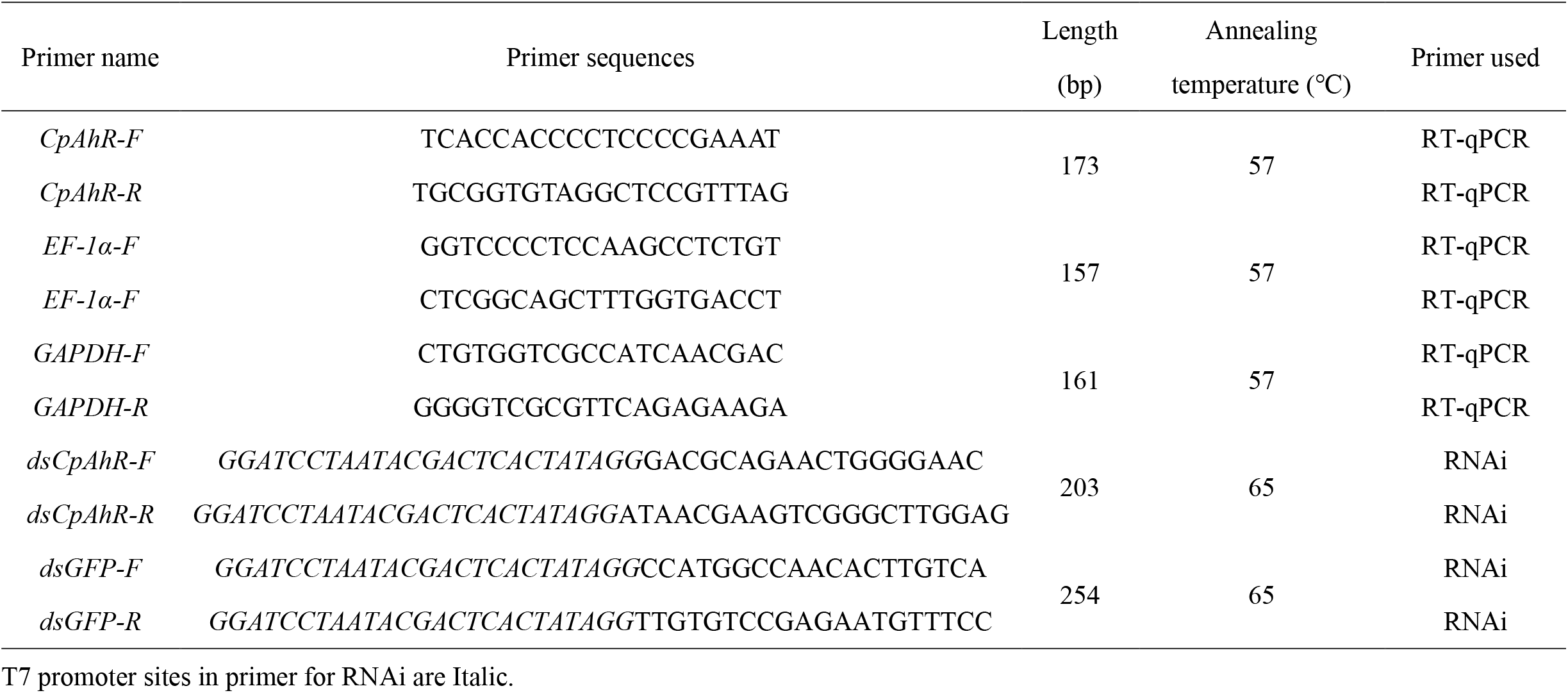
Primers used in this study.

### 2.4. Identification, sequence, and phylogenetic analysis of CpAhR

The amino acid sequences of the *AhR* gene from *Bombyx mori, Plutella xylostella, Spodoptera litura, Ostrinia furnacalis*, and *Helicoverpa armigera* (Table S1) were downloaded from the National Centre for Biotechnology Information (NCBI: www.ncbi.nlm.nih.gov/) and used as query sequences for searches of TBLASTN (e-value <10^− 5^) in the *C. pomonella* transcriptome database, which was downloaded in the genome database from InsectBase (http://v2.insect-genome.com/). The conserved protein domains of *AhR* genes were predicted using the NCBI conserved domain database (CDD) (https://www.ncbi.nlm.nih.gov/cdd). The ExPASy Proteomics Server (https://web.expasy.org/compute_pi/) was an online website for predicting molecular weight (Mw) and isoelectric point (pI). Amino acid sequences (Table S1) were aligned by ClustalW and the phylogenetic tree was constructed by the neighbor-joining method with 1000 bootstrap replicates using MEGA X software. Sequence similarity was determined by aligning with DNAMAN 5.2.2 (LynnonBioSoft, PointeClaire, Canada).

### 2.5. RNA interference (RNAi) of CpAhR

The synthesis of double-stranded RNA (dsRNA) was followed by the manufacturer’s instructions of the T7 RiboMAXTM Express RNAi System (Promega, USA). Briefly, the cDNA templates were amplified by PCR using gene-specific primers conjugated with a T7 promoter site (Table 1). The PCR amplicons were recovered and purified for the synthesis of dsRNA, and the final product was diluted to 2000 ng μL^− 1^. Before injection, test fourth-instar larvae were placed on ice for 5 min to make them faint, and then quickly transferred to the culture dish for injection. A sample of 1 µL dsRNA was injected into the 3 -5 segment at the end of the abdomen of each larva, and an equal amount of dsGFP was injected as the control. At least five larvae were injected into each group and a total of three replicates were performed. The injected samples were collected within 48 h and immediately treated with liquid nitrogen and stored at - 80 ℃.

### 2.6. Enzymatic assay

The extraction of total GST enzyme and enzymatic activity was determined according to the method of Rodríguez et al [31] with minor modifications. GST enzyme preparation was added to microplate wells containing CDNB (30 mM), and GSH (100 mM), and supplemented to 200 μL with 50 mM sodium phosphate buffer (pH 7.2). The absorbance value change of the mixture at 340 nm for two minutes was recorded and GST activity was calculated using the extinction coefficient (ε, 9.6 mM^-1^ cm^-1^).

The extraction of the total P450 enzyme and enzymatic activity was determined according to the method of Tang et al [32] with minor modifications. The extracted P450 enzyme solution was added to a centrifuge tube containing substrate 7-ethoxy-coumarin (2 mM), NADPH (10 mM), and supplemented to 200 μL with 100 mM sodium phosphate buffer (pH 7.8). After incubating the mixture at 30 °C for 10 min, 60 μL trichloroacetic acid (15%) was added to terminate the reaction. The mixture was centrifuged again at 4 °C for 10 min, and 200 μL supernatant was transferred to opaque microplate wells containing 90 μL glycine/sodium hydroxide (1.6 mM, pH=10.5). The absorption value of the mixture was recorded at the excitation wavelength of 358 nm and the emission wavelength of 456 nm, and P450 enzyme activity was assessed by measuring the amount of 7-hydroxycoumarin (ECOD) produced within 10 minutes using. After the determination, the standard curve was established according to the absorbance value of the 7-hydroxycoumarin standard of different concentrations.

### 2.7. Determination of toxicity to lambda-cyhalothrin

According to the results of the pre-experiment, the appropriate dosage of the agonist was selected to verify the increased tolerance to *lambda*-cyhalothrin. The mother liquor of *lambda*-cyhalothrin was prepared with acetone and diluted to the desired concentration (LD_50_) according to the result of Wang et al [5]. In brief, BNF was dissolved in DMSO, and then added to the artificial diet and stirred evenly. The same volume of DMSO was added as a control. The artificial feed mixed with BNF was cut into blocks of appropriate size and separately installed into the small tubes. The fourth instar larvae after starvation treatment were transferred to the tube to feed for 48 h and then treated with *lambda*-cyhalothrin of LD_50_ dose to record the mortality. Similarly, the larvae after injection of dsRNA were titrated with an LD_50_ dose of *lambda*-cyhalothrin to determine the tolerance to insecticide. One μL of insecticide solution was dropped on the pronotum of each larval individual, and the mortality was recorded after exposure. The individuals were considered dead if there is no response after gently touching them with a small brush. Each treatment was replicated at least three times with each replication containing 15 larvae.

### 2.8. Data analysis

Differences in expression levels were analyzed using one-way analysis of variance (ANOVA) with multiple comparison Tukey’s test (*P* < 0.05) for means separation using SPSS Statistics 20 (IBM, Chicago). Student’s t-test (*, *P* < 0.05) was used for the comparison of differences between two samples in enzymatic activity and mortality. The data were expressed as the mean ± SEM values from a t least triplicate experiments and plotted using the software GraphPad Prism 5 (GraphPad Software, CA).

## 3. Results

### 3.1. Sequence and phylogenetic analysis of CpAhR

Based on the genome database of *C. pomonella* from InsectBase [33], a putative *CpAhR* gene (Accession number in NCBI: OQ102528) was identified. The putative protein weight of *CpAhR* was 43.41 kDa, and the theoretical isoelectric point was 8.19 (Table S2). To inspect the evolutionary relationships of *CpAhR* and *AhR* genes from other species (Table S1), a phylogenetic tree was constructed. The result showed that *CpAhR* is more closely related to the evolution of other Lepidoptera insects (Fig. 1). Among them, *CpAhR* has the highest homology with the *AhR* gene of *B. mori*, with an amino acid similarity of 67.42% (Table S3). Besides, the result of multiple sequence alignment of *AhR* genes of different species indicated that they share the highly conserved period-ARNT-single-minded (PER-ARNT-SIM, PAS) domains (Fig. 2), whereas the transcriptional activation domains (TAD) are not so homologous. Interestingly, the same as other Lepidoptera insects, the helix-loop-helix domain (HLH) was not found in *CpAhR* (Fig. 2).

**Fig. 1.**
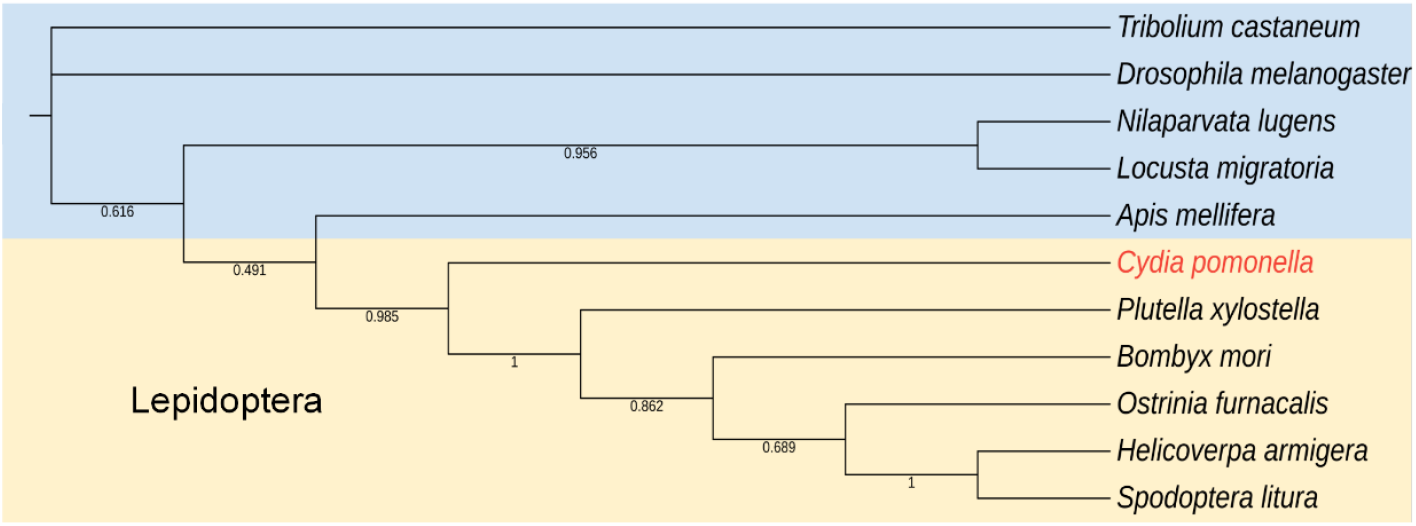
Phylogenetic analysis of *AhR* genes from *C. pomonella, B. mori, P. xylostella, S. litura, O. furnacalis, H. armigera, N. lugens, Locusta migratoria, Drosophila melanogaster, Tribolium castaneum*, and *Apis mellifera*. Numbers at the nodes of the branches represent the level of bootstrap (1000 replicates) support for each branch and the display range is set to 0.3-1. Lepidoptera insects are placed on a light yellow background, and insects other are placed on a light blue background.

**Fig. 2.**
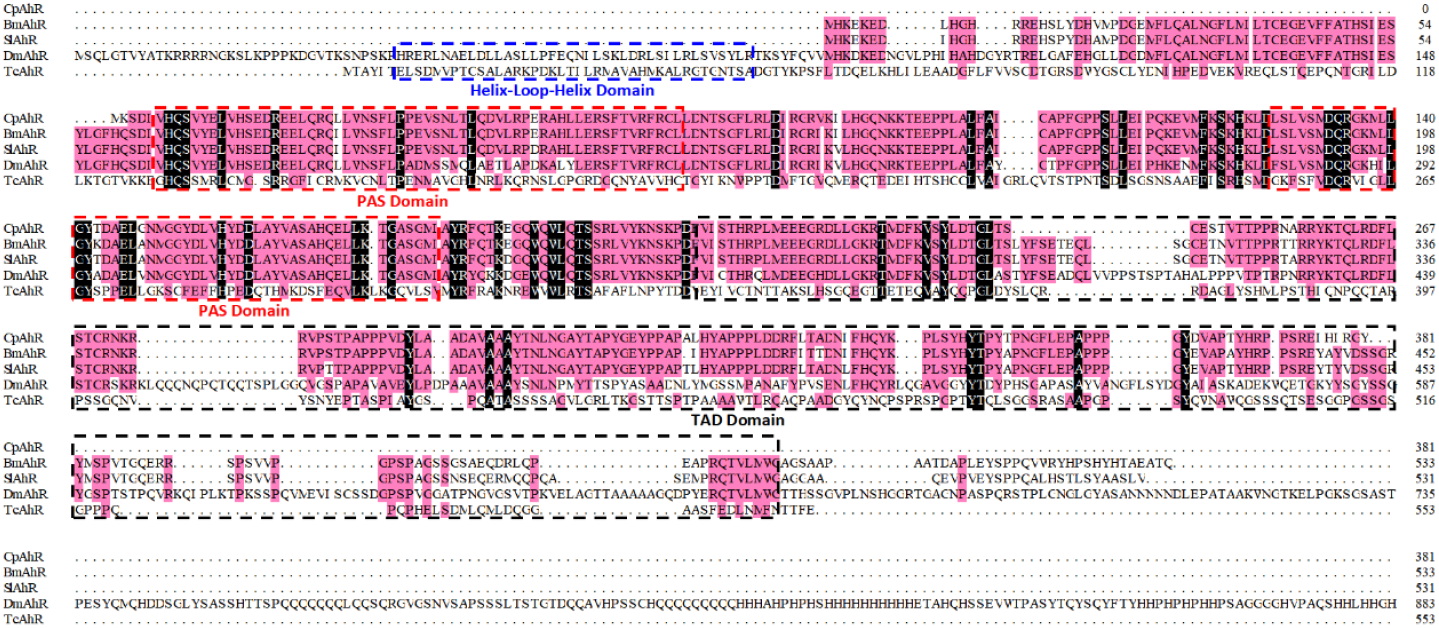
Alignment of the amino acid sequences of *AhR* genes from *C. pomonella, B. mori, S. litura, D. melanogaster*, and *T. castaneum*. The blue, red, and black dashed frames represent the helix-loop-helix domain (HLH), PER-ARNT-SIM domain (PAS), and the transcriptional activation domain (TAD), respectively.

### 3.2. CpAhR is highly expressed in the important detoxification sites

Expression patterns of *CpAhR* in developmental stages showed that the expression level of *CpAhR* in first- and fourth-instar larvae were significantly higher than those in other developmental stages (Fig. 3A). Furthermore, the expression profile of *CpAhR* in several tissues of *C. pomonella* fourth-instar larvae, including head, epidermis, fat body, midgut, and Malpighian tubules was analyzed. The relative mRNA level of *CpAhR* in Malpighian tubules was higher than in the other four tested tissues (Fig. 3B).

**Fig. 3.**
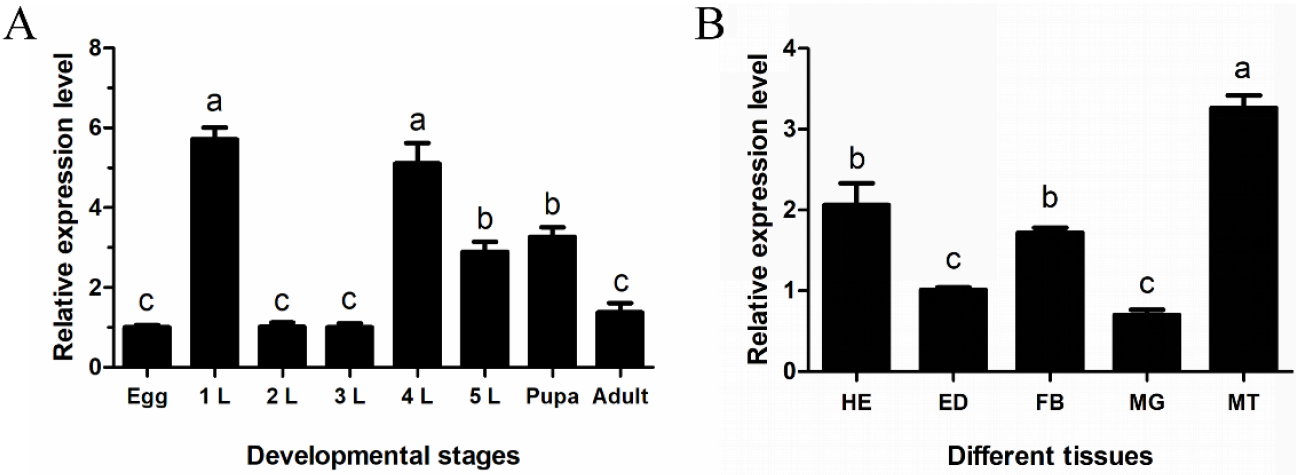
Expression levels of *CpAhR* of *C. pomonella* across different developmental stages (A) and in various tissues (B). 1L to 5L represent first- to fifth-instar larvae. HE, head; ED, epidermis; FB, fat body; MG, midgut; MT, Malpighian tubules. The results are shown as the mean ± SE. Letters on the error bars indicate significant differences analyzed by ANOVA with multiple comparisons Tukey’s test (*P* < 0.05).

### 3.3. CpAhR responses to lambda-cyhalothrin exposure

To investigate the molecular response of the transcript level of *CpAhR* to insecticide stress, fourth-instar larvae of *C. pomonella* were exposed to an LD_10_ of *lambda*-cyhalothrin. The result of RT-qPCR showed that at 6 and 12 hpe, there were no significant changes in the mRNA levels of *CpAhR* compared to the control. The gene expression level significantly increased to 5.24-fold at 24 hpe, compared to the control However, with the extension of the treatment time, the expression of *CpAhR* gradually returned to the normal level, compared with the control (Fig. 4).

**Fig. 4.**
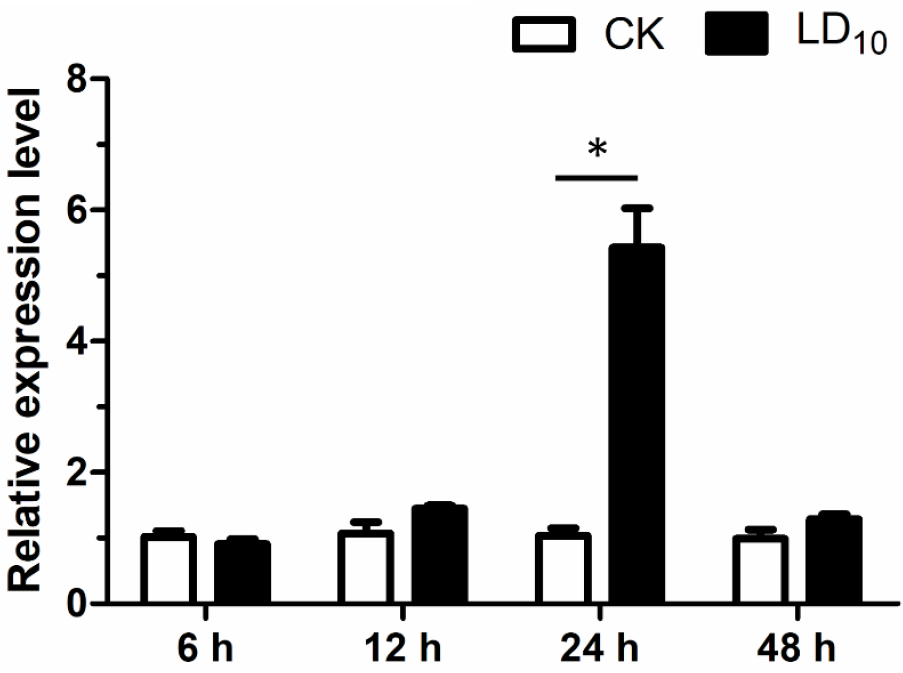
Expression levels of *CpAhR* of *C. pomonella* exposed to a sublethal dose of *lambda*-cyhalothrin. The relative expression levels at 6, 12, 24, and 48 h post insecticide exposure were calculated compared with the control treated with acetone. The results are shown as the mean ±SE. Asterisk on the error bar indicates a significant difference analyzed Student’s t-test (\**P* < 0.05).

### 3.4. CpAhR is over-expressed in lambda-cyhalothrin-resistant population

RT-qPCR showed that the expression level of *CpAhR* in the field-resistant strain (ZW_R) was 1.58-fold higher than that of its resistant regression strain, ZW_S (Fig. 5). *3.5. Silencing CpAhR reduces larval tolerance to lambda-cyhalothrin* The mRNA level of *CpAhR* decreased significantly after injection of dsCpAhR from 24 h to 48 h (Fig. 6A). Compared to the control injected with dsGFP, *CpAhR* decreased to the lowest expression level at 36 h (suppressed by 49.11 %) after injection of dsCpAhR. After the injection of dsRNA for 36 h, larvae were treated with a half-lethal dose (LD_50_) of *lambda*-cyhalothrin, and the death of each group of larvae was recorded. The results revealed that the mortality of larvae injected with dsCpAhR was significantly higher (*P*=0.0078) than that of the control as the treatment time increased to 48 h (Fig. 6B). The results of enzyme activity analysis showed that the GST enzyme activity in the larvae decreased significantly (*P*=0.0491) at 36 h after the injection of dsCpAhR (Fig. 6C), while there was no significant difference (*P*>0.05) of the P450 enzyme activity compared with control (Fig. 6D).

**Fig. 5.**
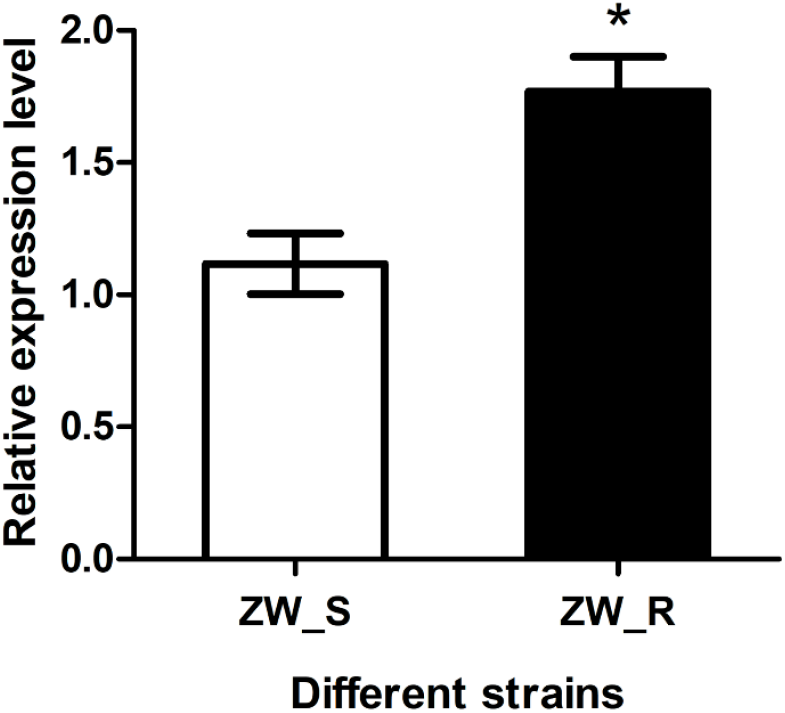
Expression profiles of *CpAhR* in the field-resistant strain of *C. pomonella*. The results are shown as the mean ± SEM. Asterisks on the error bar s indicate significant differences analyzed Student’s t-test (\**P* < 0.05).

**Fig. 6.**
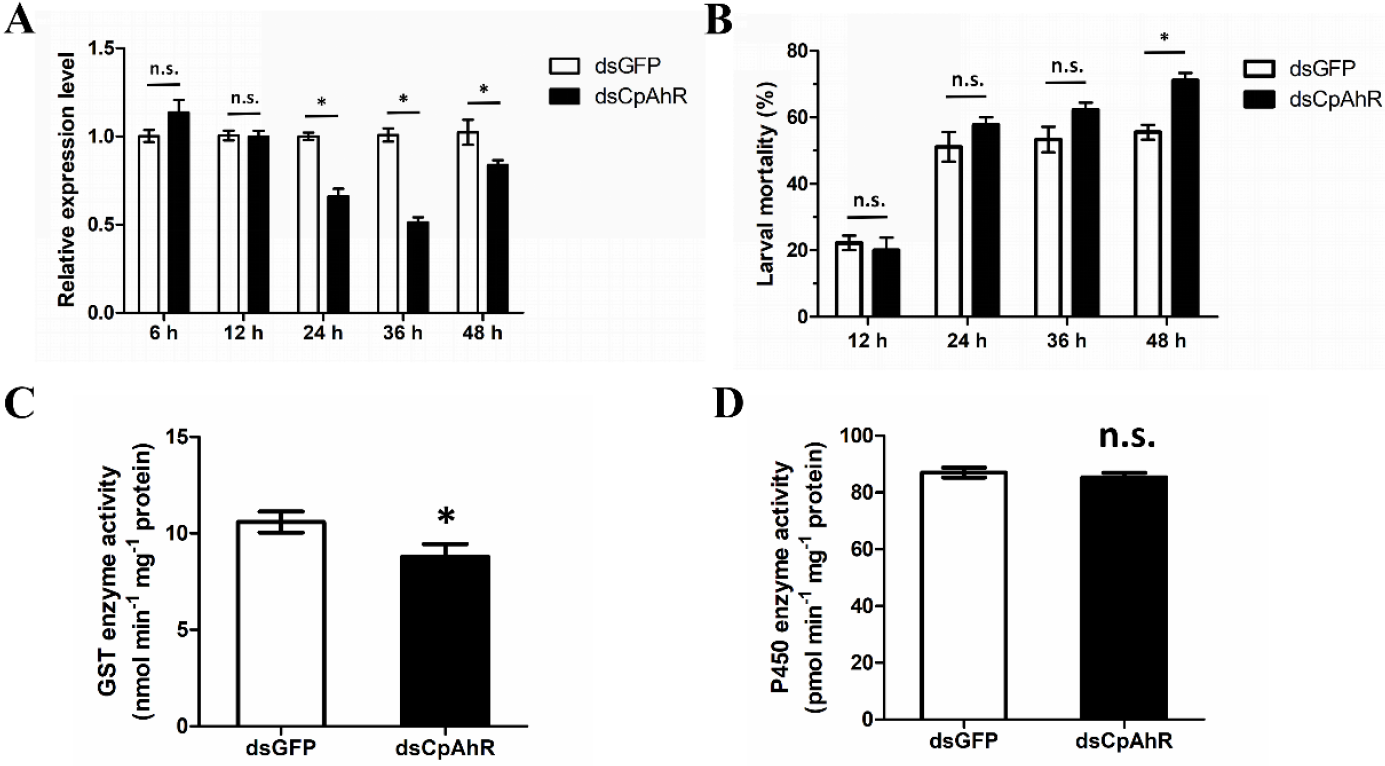
Expression levels of *CpAhR* of fourth-instar larvae injected with dsCpAhR or dsGFP at different times (A). The mortality of larvae treated with half lethal dose of *lambda*-cyhalothrin at different times after injected with dsCpAhR or dsGFP for 18 h (B). GST enzyme activity of fourth-instar larvae after injected with dsCpAhR or dsGFP for 18 h (C). P450 enzyme activity of fourth-instar larvae after injected with dsCpAhR or dsGFP for 18 h (D). The results are shown as the mean ± SEM. Asterisks on the error bars indicate significant differences analyzed Student’s t-test (\**P* < 0.05).

### 3.6. Silencing CpAhR down-regulates the expression of CpGSTe3

The previous study has proved that four GST genes, *CpGSTd1, CpGSTd3, CpGSTe3*, and *CpGSTs2* are responsible for *lambda-*cyhalothrin resistance in *C. pomonella* [8]. In this study, we detected the changes in gene expression after the injection of dsCpAhR. The result showed that compared with the control, the mRNA level of *CpGSTe3* was significantly reduced after the knockdown of *CpAhR* (Fig. 7).

**Fig. 7.**
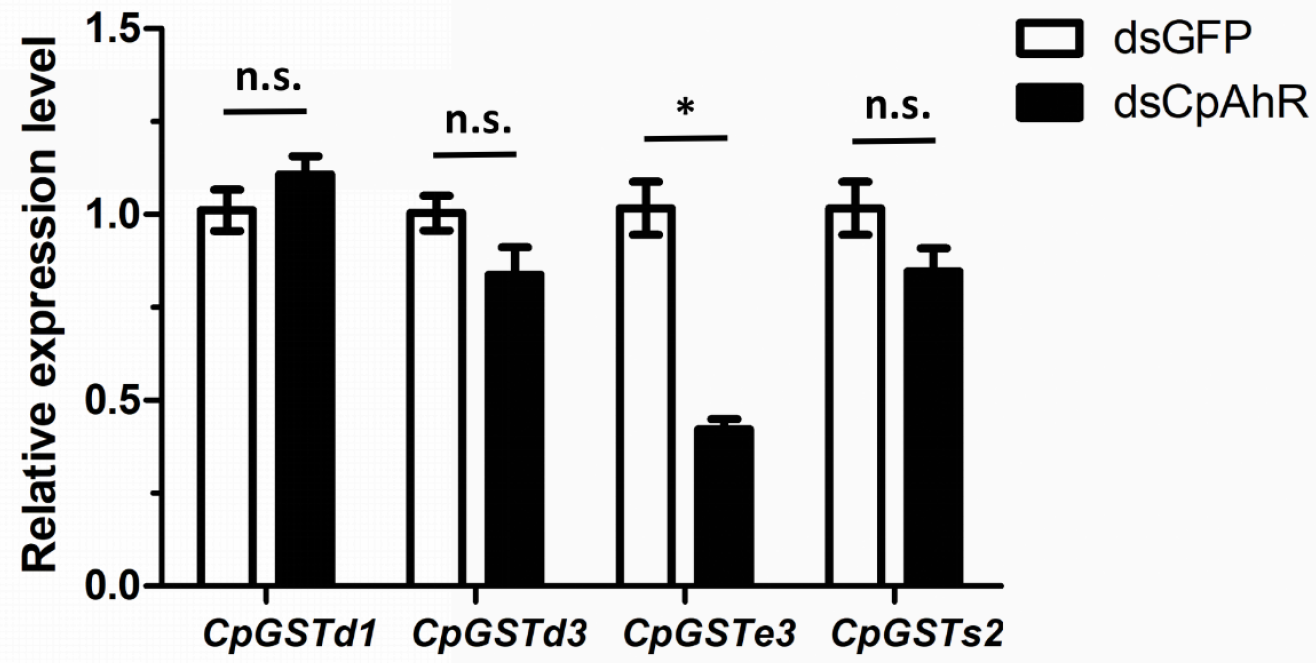
The relative expression levels of four GST genes after injection of dsRNA. The results are shown as the mean ± SEM. Asterisks on the error bars indicate significant differences analyzed Student’s t-test (\**P* < 0.05).

### 3.7. Agonist BNF activates GST activity and increases larval tolerance to lambda-cyhalothrin

Four concentrations (0, 1.0, 1.5, and 2.0 mg g^-1^) of BNF were used to feed the fourth-instar larvae to stimulate the expression of *CpAhR*. Each group of larvae did not die within 48 h after feeding on BNF with different concentrations (Fig. S1A). However, the rate of weight gain decreased significantly after taking the diet containing 2.0 mg g^-1^ of BNF, while lower concentrations of BNF did not significantly affect weight change (Fig. S1B).

The mRNA level of *CpAhR* was unaffected by 1.0 mg g^-1^ of BNF, whereas it was up-regulated by 1.98- and 1.58-fold at 1.5 and 2.0 mg g^-1^ of BNF, respectively (Fig. 8A). After feeding on BNF, an increase of tolerance was observed with the mortality of larvae treated with LD_50_ of *lambda*-cyhalothrin decreased significantly (*P*=0.0158) (Fig. 8B). Furthermore, compared with the control group, the GST and P450 enzyme activities of the larvae increased in varying degrees after being fed with different concentrations of agonists BNF (Fig. 8C and 8D). However, correlation analysis revealed that *CpAhR* expression was positively correlated with the enzyme activity GSTs (R^2^=0.9688), but not P450s (R^2^=0.2278) (Fig. S2).

**Fig. 8.**
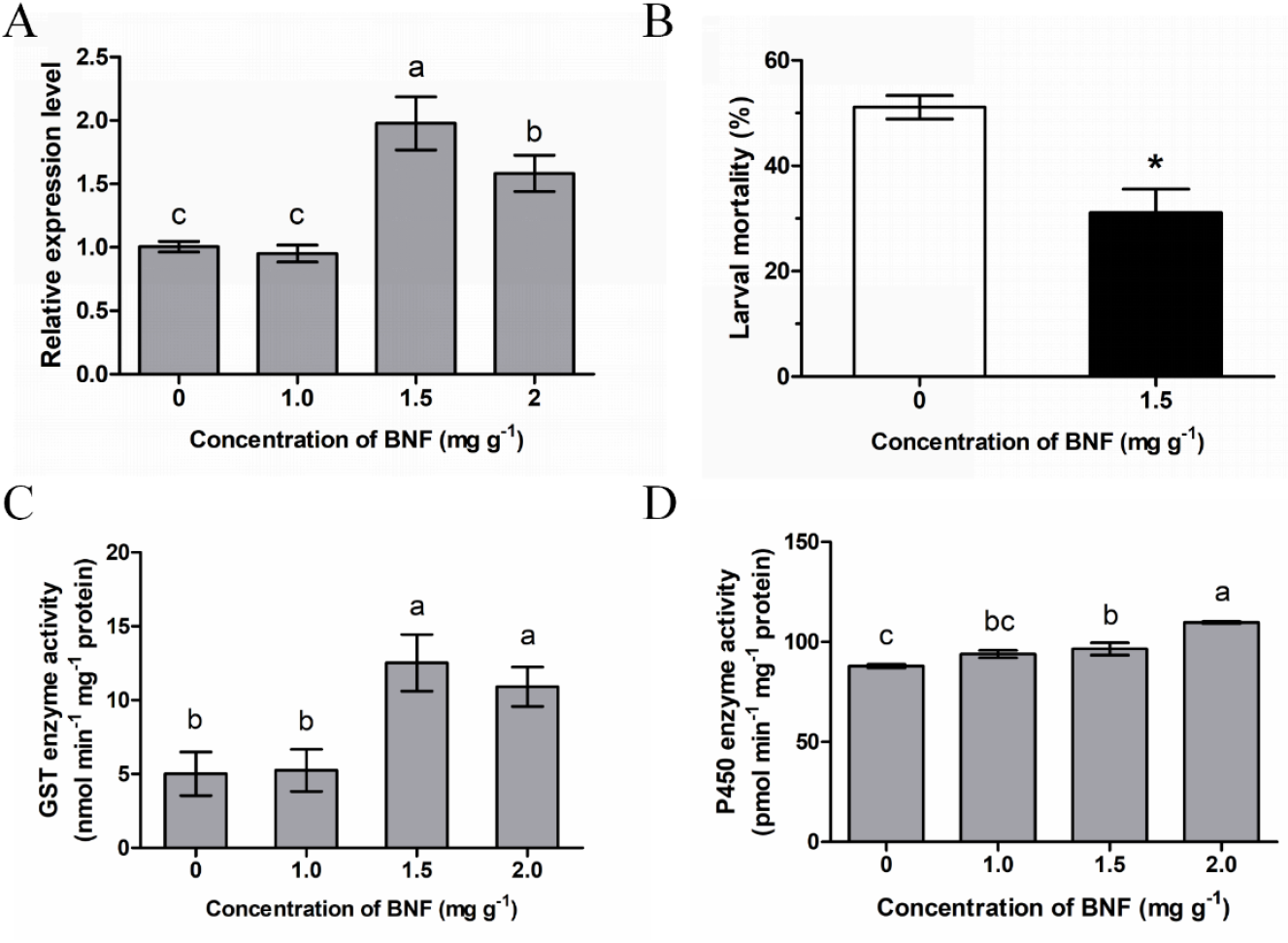
Expression levels of *CpAhR* of fourth-instar larvae after being fed with different concentrations of BNF (A). GST enzyme activity of fourth-instar larvae after being fed with different concentrations of BNF (B). P450 enzyme activity of fourth-instar larvae after fed with different concentrations of BNF (C) The mortality of larvae treated with half lethal dose of *lambda*-cyhalothrin after fed with 1.5 mg g^-1^ of BNF (D). The results are shown as the mean ± SE. Letters on the error bars indicate significant differences analyzed by ANOVA with multiple comparisons Tukey’s test (*P* < 0.05). Asterisk on the error bar indicates a significant difference analyzed Student’s t-test (\**P* < 0.05).

## 4. Discussion

*C. pomonella* is a destructive pest of pome fruits worldwide that causes a great loss of both quality and yield, and the use of chemical insecticides has been considered the most effective and direct control measure [3,34,35]. It has been demonstrated that prolonged exposure to insecticides inevitably leads to the problem of an increase in insecticide resistance, which causes almost all pests will be uncontrollable again, including *C. pomonella* [7,36,37]. Pests have developed a variety of complex regulatory mechanisms to enhance the activity of detoxification metabolic enzymes to solve the problem of persecution caused by toxin exposure [38,39]. In this study, we characterized the transcription factor *CpAhR* and confirmed that it can effectively activate the expression and enzyme activity of GSTs, but not the P450s, and thus increase the resistance of *C. pomonella* to *lambda*-cyhalothrin.

Although evolution will gradually differentiate the homologous genes in different species, the transcription factor AhR is still relatively conservative to perform specific functions [21]. We used the data in the genome from InsectBase [33] to determine the sequence and functional annotation of *CpAhR*, and also downloaded the amino acid sequences of proximate or distant insect species for phylogenetic analysis. The result showed that the *AhR* gene from Lepidoptera was well clustered on close branches and far away from other orders of insects, suggesting that there were significant differences in the evolution among different orders of insects. Besides, the helix-loop-helix domain (HLH) was not predicted in *CpAhR* as well as those of Lepidoptera insects, but different from those of other orders of insects [22,40]. It is speculated that another transcription factor ARNT that binds to AhR, contains a functional domain that combines with DNA, which leads to the degeneration of the *AhR* gene. It is probably directly caused by the evolutionary differences of Lepidoptera insects, but it is unknown whether this difference will be affected the regulation of insecticide resistance.

Although specific functional domains have been defined that cause transcription factor AhR can regulate the transcriptional activation of downstream genes, evolutionary differences by pluralistic environments will inevitably affect functional differences, which requires comprehensive analysis. Thus, due to the feeding characteristic of insects and the differences in the function of different tissues or organs *in vivo* affecting the specific expression of genes, it’s general to study the function of detoxification metabolism gene based on the spatiotemporal specific expression [41-43]. Our RT-qPCR analysis suggested that the mRNA levels of *CpAhR* were significantly high in the first- and fourth-instar larval stages, which are the initial stage and overeating stage of food intake during the life cycle of *C. pomonella*, respectively. That is because the activation of the *AhR* gene expression will positively respond to the exogenous substances contained in food [44]. Moreover, transcripts of *CpAhR* were found in all tested tissues and it was expressed to the greatest extent in the Malpighian tubules, followed by the head, fat body, epidermis, and midgut. High expression levels in Malpighian tubules and fat body may be because these two are important places for absorbing and excreting exogenous substances [8], whereas the high expression of a transcription factor that can regulate detoxification gene in the head likely aid in reducing the concentration of toxicant and mitigate harmful effects in nerve cells or tissues [45]. Although we have roughly inferred the role of *CpAhR* in detoxification metabolism based on the results of spatiotemporal expression, the specific function needs further study.

It has been proved that exposure to insecticides will induce the expression and enzyme activities of GSTs and P450s in insects [8,46], but the expression response of transcription factors to insecticide is less studied. In the present study, the relative expression level of *CpAhR* did not change significantly in the early stage of exposure to *lambda*-cyhalothrin, then increased to 5.24-fold at 24 h, and finally decreased. The such result indicated that it takes a period for the regulation pathway of *CpAhR* as the regulation center to react after *lambda*-cyhalothrin treatment, while the subsequent decline of transcription level of *CpAhR* may be due to the fulfillment of the storage stage to downstream genes. Furthermore, *CpAhR* was up-regulated in a field-*lambda*-cyhalothrin resistant strain (ZW_R), compared with its resistant regression strain (ZW_S) with the same genetic background, indicating that *CpAhR* could be contributing to *lambda*-cyhalothrin resistance through overexpression. The above results are similar to previous reports on other species. Under the long-term selection of imidacloprid pressure, *Sitobion miscanthi* gradually developed resistance to imidacloprid, and the expression level of *AhR* gene increased significantly [47]. Similarly, in *T. castaneum*, the mRNA expression level of *TcCncC* was significantly up-regulated at 24 h and 60 h after eugenol treatment [48]. That is, xenobiotics such as insecticides or plant secondary metabolites can not only induce the expression of detoxification genes, but also the transcription factors, thus participating in the detoxification regulation network and increasing the tolerance of insects to toxins. Meanwhile, insecticide stress has a positive correlation with the expression of transcription factors, which can be used as a shred of strong evidence of whether they participate in the regulation of insecticide resistance.

The low RNAi efficiency of Lepidoptera insects is due to multiple factors [49], but it doesn’t affect that it is still regarded as one of the most effective ways to verify gene function [50]. In this study, although the mRNA level of *CpAhR* decreased significantly after injection of dsCpAhR from 24 h to 48 h, the highest interference efficiency is only close to 50% and the duration was not very long. Therefore, we selected the time point (36 h) with the highest interference efficiency to determine the change of tolerance to *lambda*-cyhalothrin, and the result showed that a significant difference in larval mortality appeared after 48 h of LD_50_ *lambda*-cyhalothrin treatment, which implies that *CpAhR* is involved in the regulation of tolerance to *lambda*-cyhalothrin. Furthermore, we found that after injection of dsCpAhR, the enzyme activity of GSTs decreased significantly, whereas the activity of P450s did not change significantly, indicating that *CpAhR* enhances the tolerance to *lambda*-cyhalothrin by increasing the enzyme activity of GSTs. We further found that the mRNA level of *CpGSTe3* was significantly reduced after injection of dsCpAhR, implying that *CpGSTe3* may be regulated by *CpAhR* and play an important role in resistance to *lambda*-cyhalothrin.

BNF is a potent AhR agonist and it induces phase I enzymes, such as CYP1A1/2, and phase II enzymes inducer, such as UDP glucuronosyltransferases (UDP-GTs) and glutathione S-transferase (GST) [28,51]. However, research on AhR agonist BNF has tended to focus on partial mammals and aquatic animals, with basically no attention to insects [29]. In this study, BNF was used as an agonist for the first time to evaluate its role in the activation of AhR in insects, and the result showed that fed larvae with artificial feed mixed with 1.5 mg g^-1^ of BNF could effectively increase the expression level of *CpAhR* as well as the activity of GSTs, indicating that a close regulatory relationship ad suggests that BNF could be a potent agonist for activation of AhR of insects.

Currently, most studies have shown that *AhR* regulates P450 genes and thus confers insecticide resistance, and only a few showed that *AhR* could also regulate GST genes in insects. For example, the *AhR* gene was involved in the regulation of the xenobiotic tolerance-related cytochrome P450 *CYP6DA2* in *Aphis gossypii* [40]. AhR/ARNT regulated the expression of *CYP6CY3* and *CYP6CY4* cooperatively, conferring the nicotine adaption of *M. persicae* to tobacco [22]. In *S. miscanthi*, AhR/ARNT regulated *CYP307A2* conferring resistance to two neonicotinoid insecticides [47]. Unlike, the expression of multiple GSTs was co-regulated by transcription factors CncC/Maf and AhR/ARNT, conferring resistance to chlorpyrifos and cypermethrin in *Spodoptera exigua* [52]. Interestingly, in this study, we found that *CpAhR* was involved in the regulation of GSTs, but not P450s in *C. pomonella*. Down-regulation of *CpAhR* expression via RNAi decreased the activity of GST enzyme and enhanced the larvae tolerance to *lambda*-cyhalothrin, while up-regulation of *CpAhR* expression via agonist BNF increased the activity of GSTs and reduced the larvae tolerance to *lambda*-cyhalothrin, which reveals that *CpAhR* is involved in the regulation of detoxification enzyme GSTs and further leads to resistance to *lambda*-cyhalothrin in *C. pomonella* (Fig. 9). Our finding provides helpful theoretical support for future research to verify the function of the transcription factor AhR and more importantly enriches the theory of insecticide resistance regulation in insects.

**Fig. 9.**
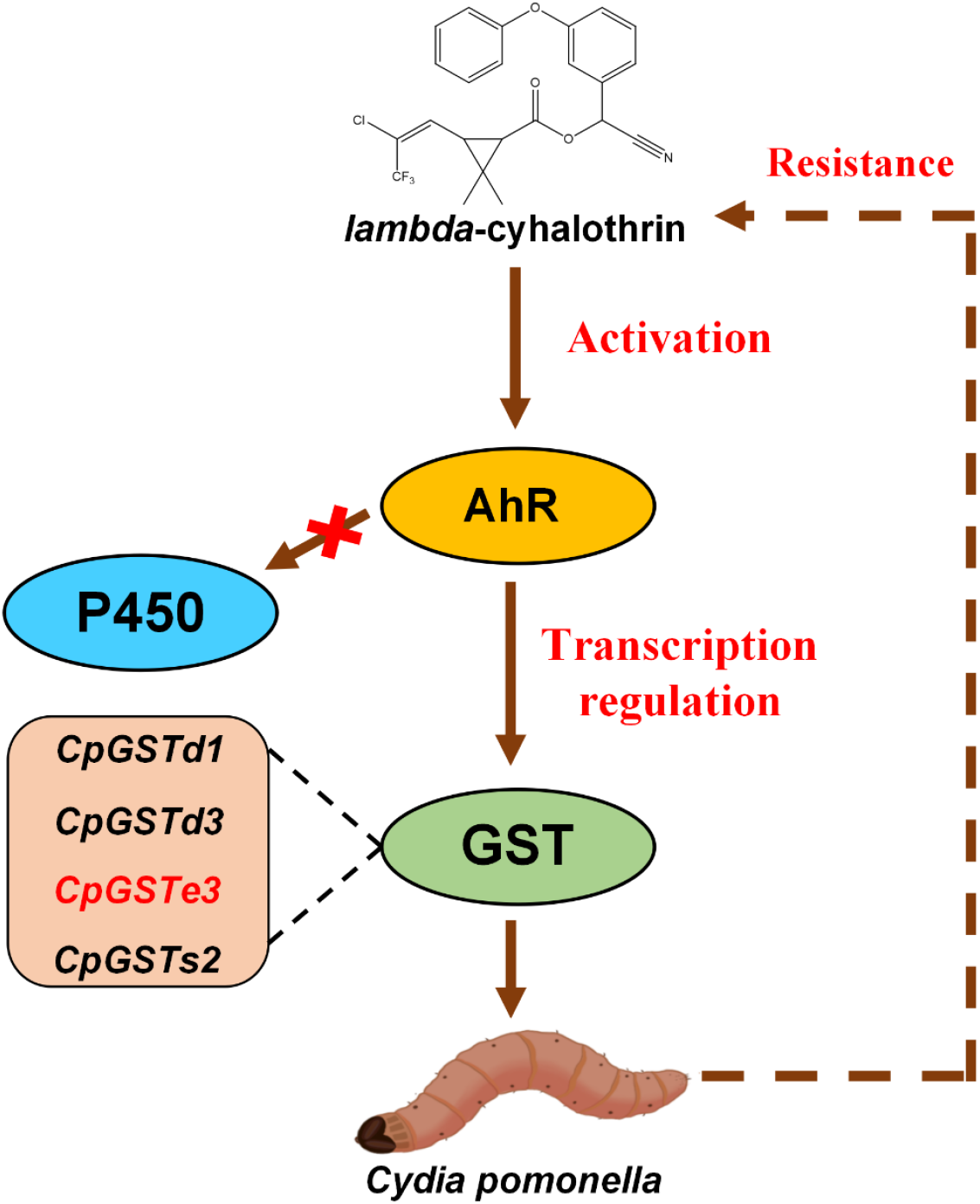
A working model of the regulatory mechanism of *CpAhR* for *lambda*-cyhalothrin resistance.

### CRediT authorship contribution statement

Xue-Qing Yang conceived and designed this study. Chao Hu and Yu-Xi Liu performed the experiments. Chao Hu collated and analyzed the data, designed the graphs, and wrote the first draft of the manuscript. Shi-Pan Zhang and Ya-Qi Wang provided several test materials and samples. Chao Hu and Yu-Xi Liu participated in the manuscript supplement and modification. Xue-Qing Yang checked and revised the final version of the manuscript. All authors contributed to the article and approved the submitted version.

## Data availability

Data will be made available on request.

## Acknowledgements

This research was supported by the National Key R&D Program of China (2021YFD1400201), the National Natural Science Foundation of China (32272588, 31972299), and the LiaoNing Revitalization Talents Program (XLYC1907097).

## Supplementary Material

The Supplementary Material is available free of charge online. Table S1-S3 and Figure S1-S2.

