## Supplementary Table S1, Table S2, Table S3, Figure S1, Figure S2 for "Transcription factor AhR regulates glutathione S-transferases (GSTs) conferring resistance to *lambda*-cyhalothrin in *Cydia pomonella*": Supplementary Material.docx

Author running head: *C. Hu et al.*

Title running head: *AhR* regulates *lambda*-cyhalothrin resistance-related GST genes in *C. pomonella*

Table S1 Accession numbers of amino acid sequences of *AhR* genes in other species.

| Insect species | Accession number in NCBI |
| --- | --- |
| *B. mori* | XP_037867954.1 |
| P. *xylostella* | XP_048478017.1 |
| *S. litura* | XP_022815374.1 |
| *O. furnacalis* | XP_028157966.1 |
| *H. armigera* | XP_049693503.1 |
| *N. lugens* | XP_039287777.1 |
| *L.migratoria* | AVL92917.1 |
| *D.melanogaster* | NP_476748.1 |
| *T. castaneum* | KYB29569.1 |
| *A. mellifera* | XP_026298107.1 |

Table S2 The putative protein weight and the theoretical isoelectric point of *CpAhR*.

| Gene name | Accession number  in InsectBase | Accession number  In NCBI | M_w_  (kDa) | pI |
| --- | --- | --- | --- | --- |
| *CpAhR* | cpo055500 | OQ102528 | 43.41 | 8.19 |

Table S3 Percentage identities (%) of the amino acid sequence of *AhR* genes among different species.

|  | *B.mori* | *S. litura* | *P. xylostella* | *O. furnacalis* | *H. armigera* | *L. migratoria* | *D. melanogaster* | *T. castaneum* | *N. lugens* | *A. mellifera* |
| --- | --- | --- | --- | --- | --- | --- | --- | --- | --- | --- |
| C. *pomonella* | 67.42 | 62.06 | 62.23 | 66.73 | 67.32 | 34.56 | 29.52 | 13.18 | 28.87 | 32.20 |

**Fig. S1.** Survival rate of fourth-instar larvae after being fed with different concentrations of BNF (A). The rate of growth weight of fourth-instar larvae after being fed with different concentrations of BNF (B).

**Fig. S2.** The correlation analysis of *CpAhR* expression with enzyme activity of GSTs (A) and enzyme activity of P450s (B).

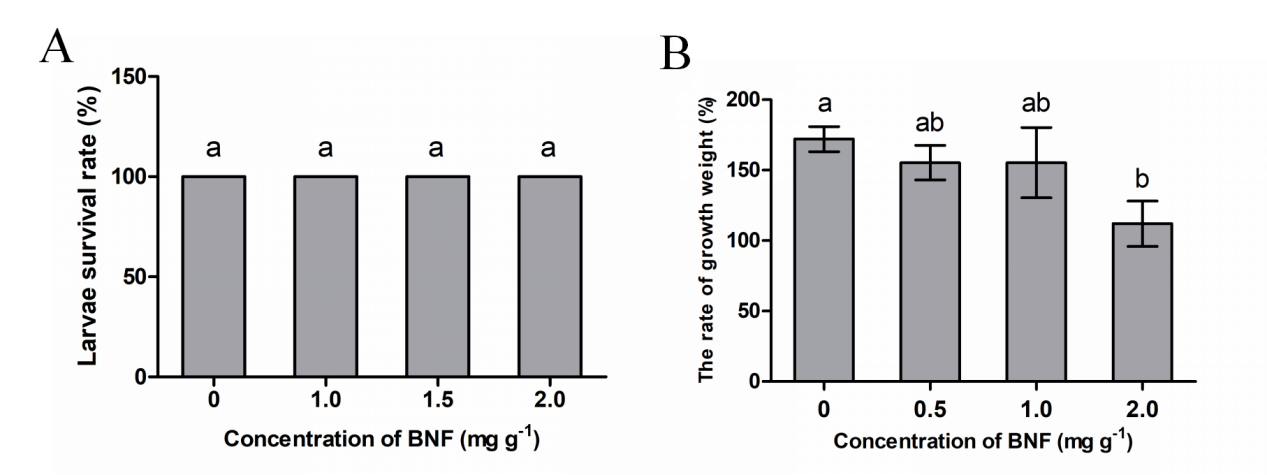

**Figure S1**

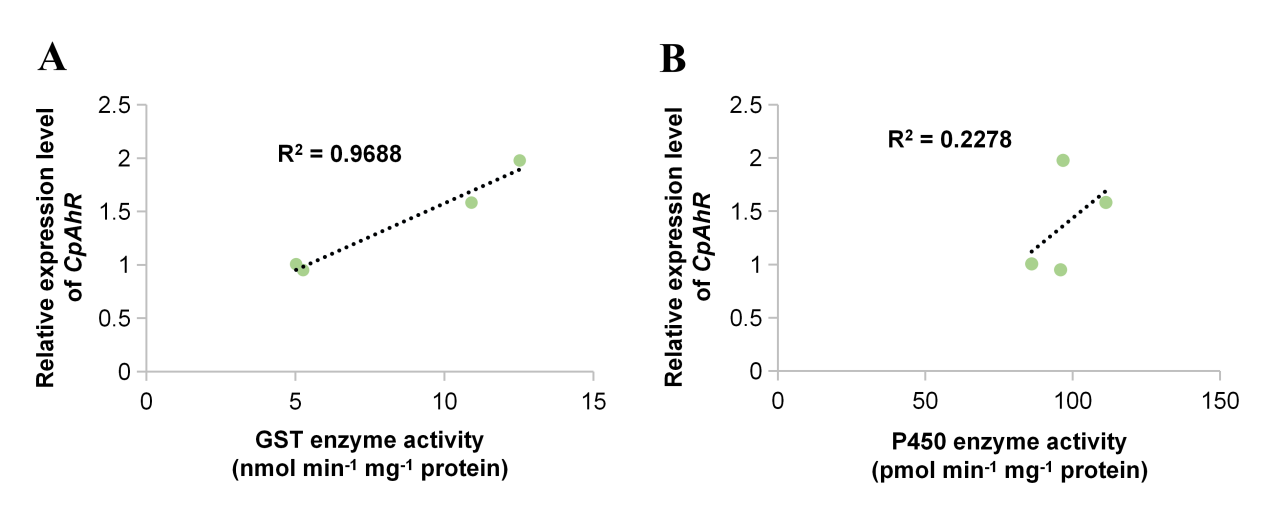

**Figure S2**
